# An eye-targeted double-RNAi screen reveals negative roles for the Archipelago ubiquitin ligase and CtBP in *Drosophila* Dpp-BMP2/4 signaling

**DOI:** 10.1101/233791

**Authors:** Nadia Eusebio, Paulo S. Pereira

## Abstract

To regulate animal development, complex networks of signaling pathways maintain the correct balance between positive and negative growth signals, ensuring that tissues achieve proper sizes and differentiation patterns. In *Drosophila*, Dpp, a member of the TGFβ family, plays two main roles during larval eye development. In the early eye primordium, Dpp promotes growth and cell survival, but later on, it switches its function to induce a developmentally-regulated cell cycle arrest in the G1 phase and neuronal photoreceptor differentiation. To advance in the identification and characterization of regulators and targets of Dpp signaling required for retinal development, we carried out an *in vivo* eye-targeted double-RNAi screen to identify *punt* (Type II TGFβ receptor) interactors. Using a set of 251 genes associated with eye development, we identified Ago, Brk, CtBP and Dad as negative regulators of the Dpp pathway. Interestingly, both Brk and Ago are negative regulators of tissue growth and Myc activity, and we show that increased tissue growth ability, by overexpression of Myc or CyclinD-Cdk4 is sufficient to partially rescue *punt*-dependent growth and photoreceptor differentiation. Furthermore, we identify a novel role of CtBP in inhibiting Dpp-dependent Mad activation by phosphorylation, downstream or in parallel to Dad, the inhibitory Smad.

## 1. Introduction

Evolutionarily conserved Transforming growth factor-β (TGFβ) signalling allows animal cells to drive developmental programmes through the regulation of cellular growth, proliferation, differentiation and morphogenesis. Its importance is reflected in the association of deregulation of this pathway with severe diseases and cancers (Massague, 2012). *Drosophila* Dpp is a ligand member of the TGFβ superfamily that signals through its type II receptors, Punt and Wit, which once activated and phosphorylated bind to type I receptors, Tkv and Sax (Restrepo et al., 2014). Together, type I and type II receptors phosphorylate the R-Smad, Mad, promoting its homodimerization and the formation of a trimeric complex with the common co-Smad Medea. This complex is then translocated to the nucleus where it controls the expression of target genes (Dijke and Hill, 2004; Rahimi and Leof, 2007). This signaling pathway is negatively regulated by the I-Smad, Dad, which competes with RSmads for receptors or Co-Smad interactions (Kamiya et al., 2008; Tsuneizumi et al., 1997). Dpp plays multiple roles in development, including the regulation of patterning and growth of the eye and wing imaginal discs (Akiyama and Gibson, 2015; Restrepo et al., 2014; Romanova-Michaelides et al., 2015). Dpp expression is required for early larval growth of the eye imaginal disc and mutations that decrease *dpp* expression also result in severely reduced adult retinas (Blackman et al., 1991; Masucci et al., 1990; St Johnston et al., 1990). Supporting these early observations, cell-autonomous activation of the Dpp pathway, through the clonal expression of constitutively-active Tkv^Q235D^ receptor was shown to increase the proliferation of eye progenitor (Firth et al., 2010). Furthermore, Tkv and Mad were shown to act cooperatively with the transcriptional coactivator Yorkie to promote retina growth (Oh and Irvine, 2011). Tissue overgrowths driven by co-expression of retinal progenitors transcription factors Hth (TALE-class homeodomain) and Tsh (zinc finger) in the eye disc causes phosphorylation and activation of Mad and depend on Dpp/BMP2 signaling for growth (Neto et al., 2016). However, the underlying mechanisms of Dpp/BMP-induced growth of the eye disc are poorly understood. Interestingly, we showed that the type II receptor Punt interacts genetically with Vito in both eye disc growth and in the onset of photoreceptor differentiation (Marinho et al., 2013). Vito is a transcriptional target of Myc and encodes an 5’ RNA exonuclease regulating rRNA and ribosome biogenesis in the nucleolus (Marinho et al., 2011). The second branch of the TGFβ signaling superfamily, the TGFβ/Activin pathway, has also been shown to be required for cell growth in the salivary glands through the control of ribosome biogenesis (Martins et al., 2017).

Retinal differentiation starts during the late L2 - early L3 larval stage within the morphogenetic furrow (MF), an epithelial indentation that advances from the posterior margin to the anterior region of the eye imaginal disc (Bessa and Casares, 2005; Fasano et al., 1991; Treisman, 2013). The progression of the MF through the eye imaginal disc, and therefore differentiation of photoreceptors, require the secretion of Hedgehog (Hh) in and behind the MF (Treisman and Heberlein, 1998). Hh controls the expression of Dpp in the MF (Greenwood and Struhl, 1999; Heberlein et al., 1995; Heberlein et al., 1993), which is necessary to switch the progenitor cell state into the precursor state allowing the initiation and progression of retinal differentiation (Bessa et al., 2002; Lopes and Casares, 2010). At this stage, Dpp has been proposed to act by repressing, at long range, transcription of *hth* that is required to maintain cells in a proliferative and undifferentiated progenitor state. Progenitor cells anterior to the furrow divide asynchronously, and Dpp also promotes G1 arrest within the furrow (Firth and Baker, 2009; Horsfield et al., 1998; Penton et al., 1997).

In this work, we studied the regulation of Dpp-BMP2/4 signaling during imaginal eye disc development. We have used an eye-targeted double RNAi screen to identify novel genetic interactions between Dpp-BMP2/4 signaling and other proteins regulating cell growth and differentiation, such as the polyubiquitin ligase component, Ago, and the transcriptional repressor CtBP. Our detailed characterization of these interactions showed that CtBP and Ago regulated eye development by different processes. Ago regulated eye disc development by promoting the critical size for eye differentiation and CtBP regulated differentiation through a negative regulation of Mad phosphorylation.

## 2. Material and Methods

### 2.1. Fly strains and genotypes

All crosses were raised at 25°C under standard conditions. Eye-targeted RNAi knockdown of *punt* was induced by crossing eyeless-Gal4 with UAS-*punt* RNAi, VDRC #37279. The overexpression of the several genes used in this work was performed using UAS-*dad*^OE^, UAS-*punt*^OE^ (a gift from Konrad Basler), UAS-*CtBP*^OE^ FlyORF #F001239, UAS-*CycB*^OE^, FlyORF #F001176, UAS-*CycB*^OE^, FlyORF #F001154, UAS-*CycD*^OE^, FlyORF #F001220, UAS-*CycD*^OE^ – *Cdk4*^0E^) (Datar et al., 2000b), UAS-*CycE*^OE^, FlyORF #F001239 and UAS-*myc*^OE^ (a gift from Filipe Josué).

### 2.2. Generation of Mosaics

Flip-out *punt* RNAi clones were generated by crossing ywhs-flp122; act>y+>Gal4 UAS-GFP females with UAS *punt* RNAi^37279^ males. Clones of cells expressing *punt* RNAi were induced at 48–72 hours after egg laying by 1 hour heat shock at 37°C. Mitotic *CtBP* mutant clones were generated by crossing ey>flip;;*CtBP*^KG07519^ FRT82B/TM6B females with M(3) Ubi GFP FRT82B/TM6B males.

### 2.3. Double-RNAi screen and genetic interaction scores

All 365 UAS-RNAi (supplementary Table S1) were obtained from VDRC, NIG-Fly stock center (http://www.shigen.nig.ac.jp/fly/nigfly/index.jsp) and Transgenic RNAi Project (TRiP) at Harvard Medical School. Eye-targeted RNAi knockdown was induced by crossing males carrying an inducible UAS-RNAi construct with eyeless-Gal4-UAS-*punt* RNAi females. All crosses were done at 25°C. The flies were examined under a stereomicroscope (Stemi 2000, Zeiss) equipped with a digital camera (Nikon Digital Sight DS-2Mv), and several representative pictures for each transgenic line were taken, if significant alterations in eye size were detected. The genetic interactions of target genes with *punt* RNAi were evaluated by comparing the phenotypes of the double RNAis versus *punt* RNAi as reference. Phenotypes were classified as lethal, sublethal if only less of 10% of the pupae hatched, small (+) retina size, medium (++) retina size if there was a significant increase in the eye size, and strong (+++) if retina size was similar to wild-type. Moreover, for the double RNAi genotypes that presented a significant increase in the retina size (+++), a supplementary evaluation was performed where adult eye size was evaluated using the following rankings: 0-25% if retinas were absent or severely reduced in size; 25-75% if retinas had moderate size reductions; and >75% if retinas had normal or nearly normal sizes.

### 2.4. Immunostaining

Immunohistochemistry of dissected eye-antennal discs was performed using standard protocols. Primary antibodies used were: rat anti-Elav 7E8A10 at 1:100 (DSHB Rat-Elav-7E8A10), rabbit anti-CtBP (kind gift of Dr. David Arnosti) at 1:5000 and rabbit anti-P-Smad1/5 41D10 at 1:100 (Cell Signaling 9516) Appropriate Alexa-Fluor conjugated secondary antibodies were from Molecular Probes. Images were obtained with the Leica SP5 confocal system and processed with Adobe Photoshop CS6.

### 2.5. Western Blot Analysis

For western blot analysis, eye imaginal discs were dissected from L3 larvae in lysis buffer (75mM HEPES pH 7.5, 1.5mM EGTA, 1.5 mM MgCl_2_, 150 mM KCl, 15% glycerol and 0.1% NP-40) containing a complete protease (Roche) and phosphatase (Sigma) inhibitor cocktails. The eye imaginal discs were homogenized with a plastic pestle. Then, the homogenized was sonified twice for 10 sec. Lysates were clarified by centrifugation for 10 min at 4°C and boiled in 1×Laemmli buffer. Protein extracts were separated by 13% SDS-PAGE and transferred to PVDF membrane. Membranes were blocked 1 h at room temperature with 5% milk in tris-buffered saline and then incubated overnight with primary antibodies at 4°C. Antibodies were diluted as follows: rabbit anti-CtBP (Dr. David Arnosti) at 1:10000 and mouse anti-tubulin β-5-1-2 (Santa Cruz Biotech) at 1:100000. Blots were detected using goat anti-rabbit and anti-mouse secondary antibodies and visualized with ECL Blotting Substrates 1:1 (Rio-Rad). A GS-800 calibrated densitometer system was used for quantitative analysis of protein levels.

### 2.6. Statistical Analysis

GraphPad Prism 5.0 was used for statistical analysis and for generating the graphical output. Statistical significance was determined using an unpaired two-tailed Student’s t-test, with a 95% confidence interval, after assessing the normality distribution of the data with the D’Agostino-Pearson normality test.

## 3. Results

### 3.1. A *Drosophila* double-RNAi combinatorial screen identifies *punt* interactors during eye development

Eye-targeted knockdown of the type-II receptor *punt* using ey-Gal4 driven expression of UAS-*punt* RNAi causes absence or very strong reduction of adult retinal tissue in the majority of animals (Figure 1; (Marinho et al., 2013; Martins et al., 2017). Additionally, 42% of ey>*punt* RNAi animals die during pupal stage. In order to identify genes that cooperate with Dpp-BMP2/4 during eye development, we performed a combinatorial double-RNAi test for modifiers of the *punt* RNAi eye phenotype. Using a set of 365 lines, we were able to study 251 different genes (Figure 1A and Table S1). The core of this gene set was described previously (Marinho et al., 2013) and contain genes functionally classified as being involved in eye development, cell cycle, transcription, or translation. We also included several members of signaling pathways important for growth and patterning during eye development, such as Hedgehog (Hh), Notch and Wingless (Wg).

**Figure 1.**
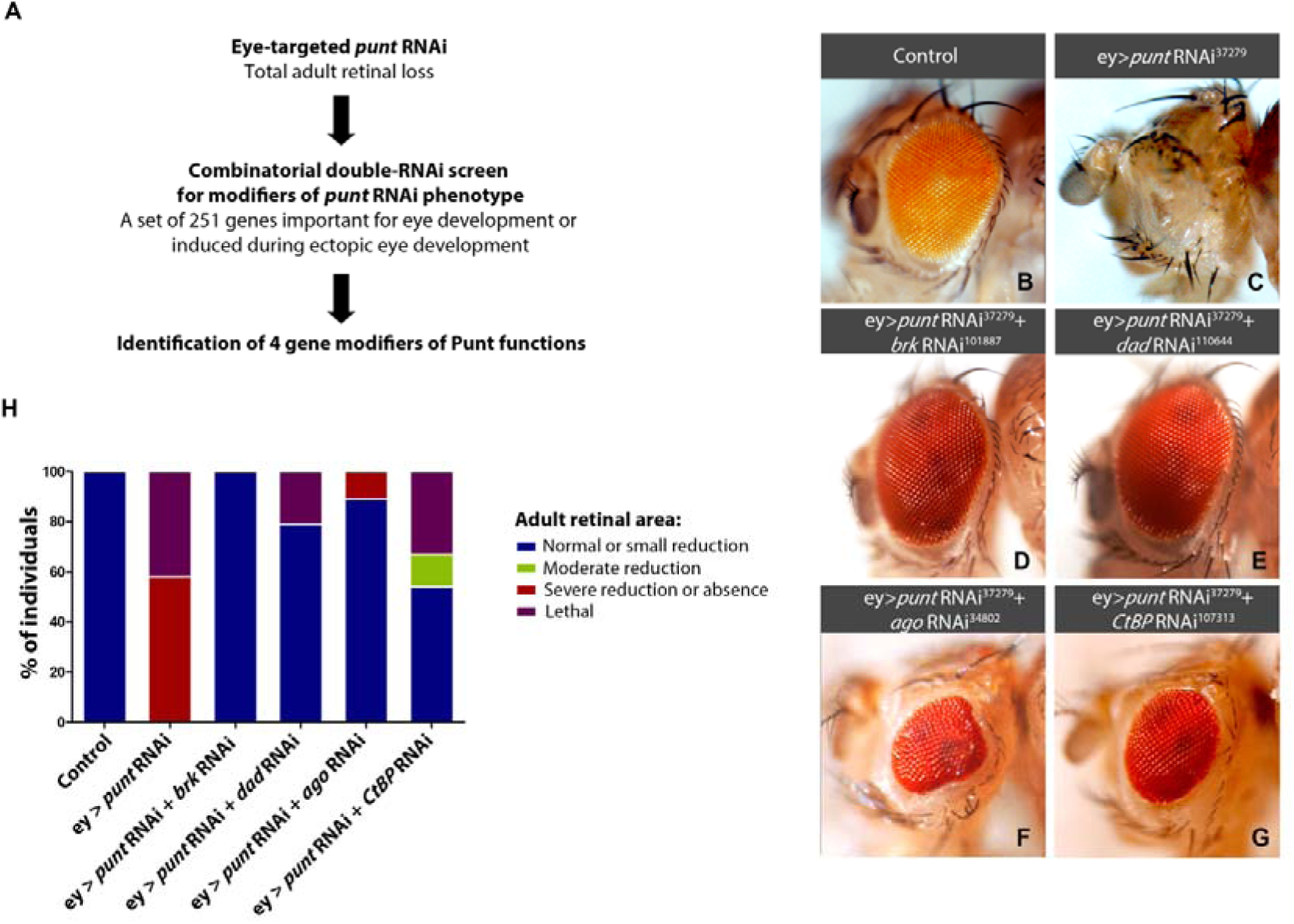
Identification of four gene modifiers of Punt loss-of-function phenotype. (A) Eye-targeted double-RNAi screen approach for identification of genes functioning with *punt* during eye development. (B–G) Representative images of the adult eye phenotypes of the indicated genotypes. ey>*punt* RNAi^37279^ shows a very strong eye phenotype without retinal formation (C). However, the differentiation failure phenotype of ey>*punt* RNAi^37279^ is rescued by *brk* RNAi^101887^ (D), *dad* RNAi^110644^ (E), *ago* RNAi^34802^ (F) and *CtBP* RNAi^107313^ (G). (H) Percentage of individuals of the indicated genotypes presenting normal adult retinal area or small reduction of adult retinal area (+++), moderate reduction of adult retinal area (++), severe reduction or absence of adult retinal area (+ or no retina) and lethality in pupa (n=43−96).

From the 365 RNAi lines tested in combination with ey>*punt* RNAi, 66 lines induced some rescue of the ey>*punt* RNAi absent eye phenotype (Table S1). Within the remaining 299 lines, 232 lines did not modify the ey> *punt* RNAi phenotype and 67 lines enhanced it, having a lethal (58 lines) or sublethal (9 lines) phenotype. From the 66 lines that rescued the eye phenotype of *punt* knockdown, only the knockdown of five candidate genes was able to strongly rescue the phenotype (>75% of the normal retina size). Interestingly, we observed that knocking down either Dad, the I-Smad, or Brk (a transcriptional repressor of Dpp targets) rescued eye development when co-expressed with ey>*punt* RNAi (Figure1D and 1E) (Bray, 1999; Kamiya et al., 2008). These results support the potential of the eye double-RNAi screen to identify Dpp regulators. The other three genes that presented a strong interaction with *punt* were *ago*, *CtBP* (Figure 1F and 1G) and *ND75* (data not show). To overcome potential off-target effects in our screen (Dietzl et al., 2007), we tested further available RNAi lines for *punt* interactors, obtaining very similar results to all genotypes (Figure S1), with the exception of *ND75* (data not shown). Flies expressing RNAis against *dad*, *brk*, *ago*, and *CtBP* did not show significant defects in adult eyes (Figure S2).

### 3.2. CtBP and Ago genetically interact with the Dpp pathway during eye development

Our genetic screen for *punt* interactors during eye development identified a strong genetic interaction with *CtBP* and *ago*. Photoreceptor differentiation was not observed in ey> *punt* RNAi eye imaginal discs (Figure 2A and 2B). However, differentiation was strongly rescued if *CtBP* RNAi or *ago* RNAi were co-expressed, even though a delay of the MF progression at the margins was observed (Figure 2C and 2D). This phenotype is also expressed by partial Dpp loss-of-function genotypes, including the hypomorph *dpp^blk^* mutant (Marinho et al., 2013). Additionally, co-depletion of *punt* together with *ago* resulted in eye discs with increased tissue growth compared to *punt* depletion alone (Figure 2D).

**Figure 2.**
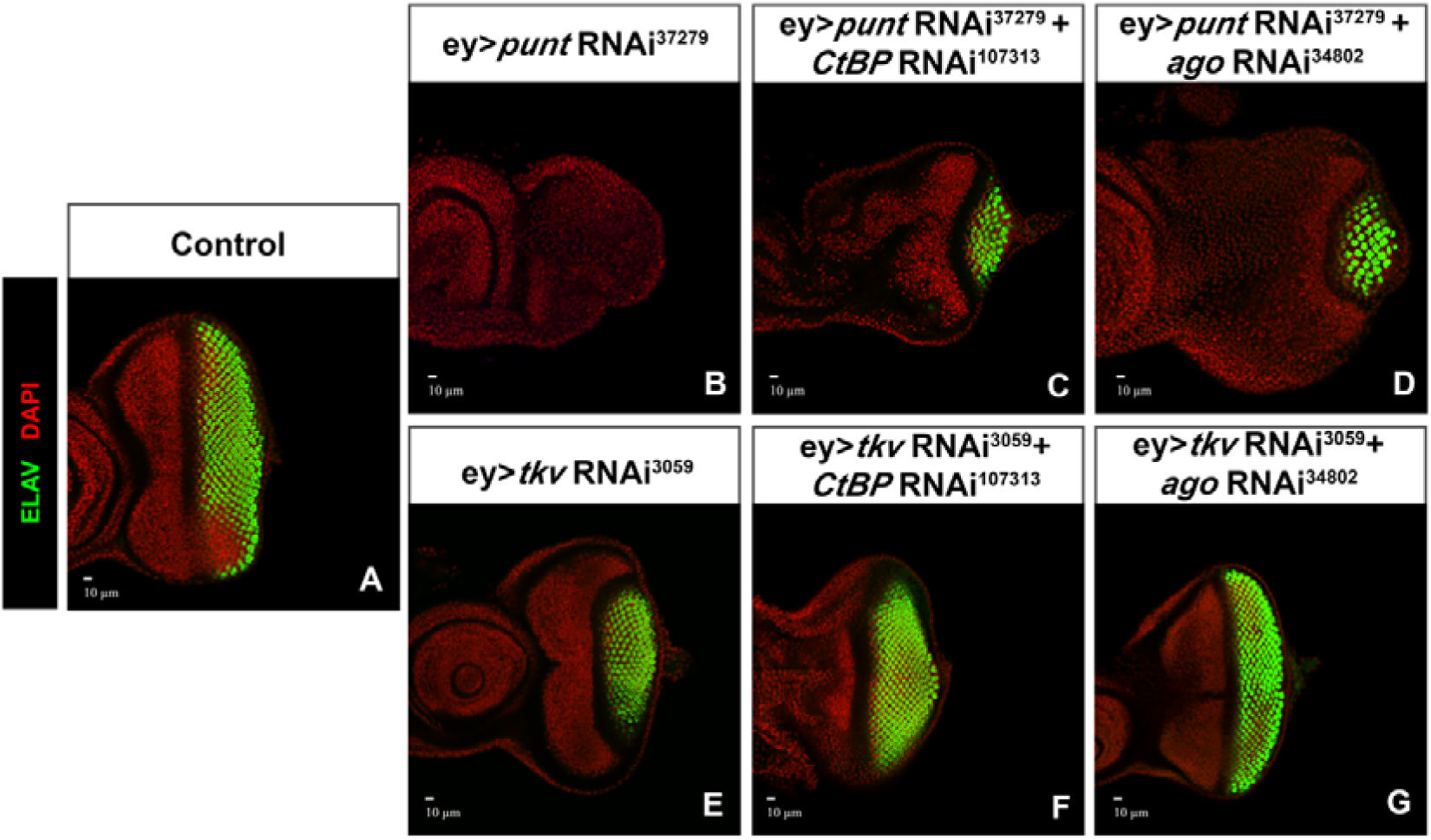
CtBP and Ago genetically interact with the Dpp-BMP2/4 signaling. Downregulation of Dpp signaling using ey>*punt* RNAi^37279^ blocks photoreceptor differentiation (A, B). (C-D) The combinatorial RNAi downregulation of *punt* together with *CtBP* or *ago* partially rescues photoreceptor differentiation (ey>*punt* RNAi^37279^+ *CtBP* RNAi^107313^; ey>*punt* RNAi^37279^+ago RNAi^34802^). (E) The downregulation of Dpp signaling using a RNAi for *tkv* leads to an impairment of differentiation progression at eye imaginal disc margins (ey>*tkv* RNAi^3059^) (F, G). The combinatorial downregulation of *tkv* together with *CtBP* or *ago* by RNAi (ey>*tkv* RNAi^3059^+ *CtBP* RNAi^107313^; ey> *tkv* RNAi^3059^+*ago* RNAi^34802^) rescues the differentiation delay at the margins induced by *tkv* RNAi. Eye discs were stained with DAPI (DNA, red) and anti-ELAV (photoreceptors, green). Scale bars correspond to 10 μm.

The canonical TGFβ signaling pathway can be divided into two main branches, BMP2/4 and Activin, which are activated by specific ligands but share a requirement for the Punt type-II receptor. Therefore, for further validation we tested the interactions of *CtBP* and *ago* with *tkv*, the Dpp BMP2/4 dedicated type-I receptor. Eye-targeted *tkv* knockdown delays progression of photoreceptor differentiation at the eye imaginal disc margins (Marinho et al., 2013), which mimicked the hypomorphic *dpp* mutant phenotype (Chanut and Heberlein, 1997) (Figure 2E and 2A), and a slight reduction in adult eye size (Marinho et al., 2013) (Figure S3). Importantly, these phenotypes were rescued by co-expressing *CtBP* RNAi and *ago* RNAi (Figure 2F and 2G).

### 3.3. Loss of Archipelago function and conditions that stimulate tissue growth rescue initiation of photoreceptor differentiation caused by Punt depletion

We observed that knocking down Ago function is sufficient for initiation and progression of photoreceptor differentiation in ey> *punt* RNAi eye discs (Figure 2D and 3A). Ago is an F-box protein that acts as the substrate-receptor component of a Skp/Cullin/F-box (SCF) E3 ubiquitin ligase (SCF-Ago) that targets Myc and CycE for degradation (Moberg et al., 2001; Moberg et al., 2004; Welcker and Clurman, 2008). Loss of Ago in imaginal discs causes an accumulation of Cyclin E and Myc, which drive cell growth and proliferation (Moberg et al., 2001; Moberg et al., 2004). We observed that rescue of *punt* RNAi eye phenotype by *ago* RNAi indeed correlated with increased Myc protein expression (Figure S4). Next, we tested the hypothesis that knockdown of *ago* rescues Dpp BMP2-4 signaling through the detected Myc upregulation, given the previously described genetic interaction between overexpression of Myc and Punt in eye growth and differentiation (Martins et al., 2017). Indeed that was the case, as overexpression of Myc was also sufficient for a partial rescue of differentiation in all the eye discs and adults that we observed (Figure 3B, 3F, 3I). Importantly, this rescue was specific as Myc could not rescue the small retina size caused by expression of a dominant-negative Jak/STAT ligand (Dome^Δcyt^) (Tsai and Sun, 2004) (Figure S5). As Ago not only targets Myc for degradation but also CycE (Koepp et al., 2001; Moberg et al., 2001), we also tested if the overexpression of CycE (*CycE*^OE^) rescued *punt* RNAi phenotype (Figure 3C, 3G and 3I). Interestingly, both the overexpression of CycE or of the CycD-Cdk4 (Datar et al., 2000a) led to a weaker, but significant, rescue of *punt* RNAi phenotype while overexpression of the CycB or CycA failed to do so (Figure 3). Taking all together, these results suggest that multiple condition that lead to growth stimulation could be sufficient to promote the onset and progression of the MF, enabling photoreceptor differentiation in eye discs with attenuated or compromised levels of Dpp signaling.

**Figure 3.**
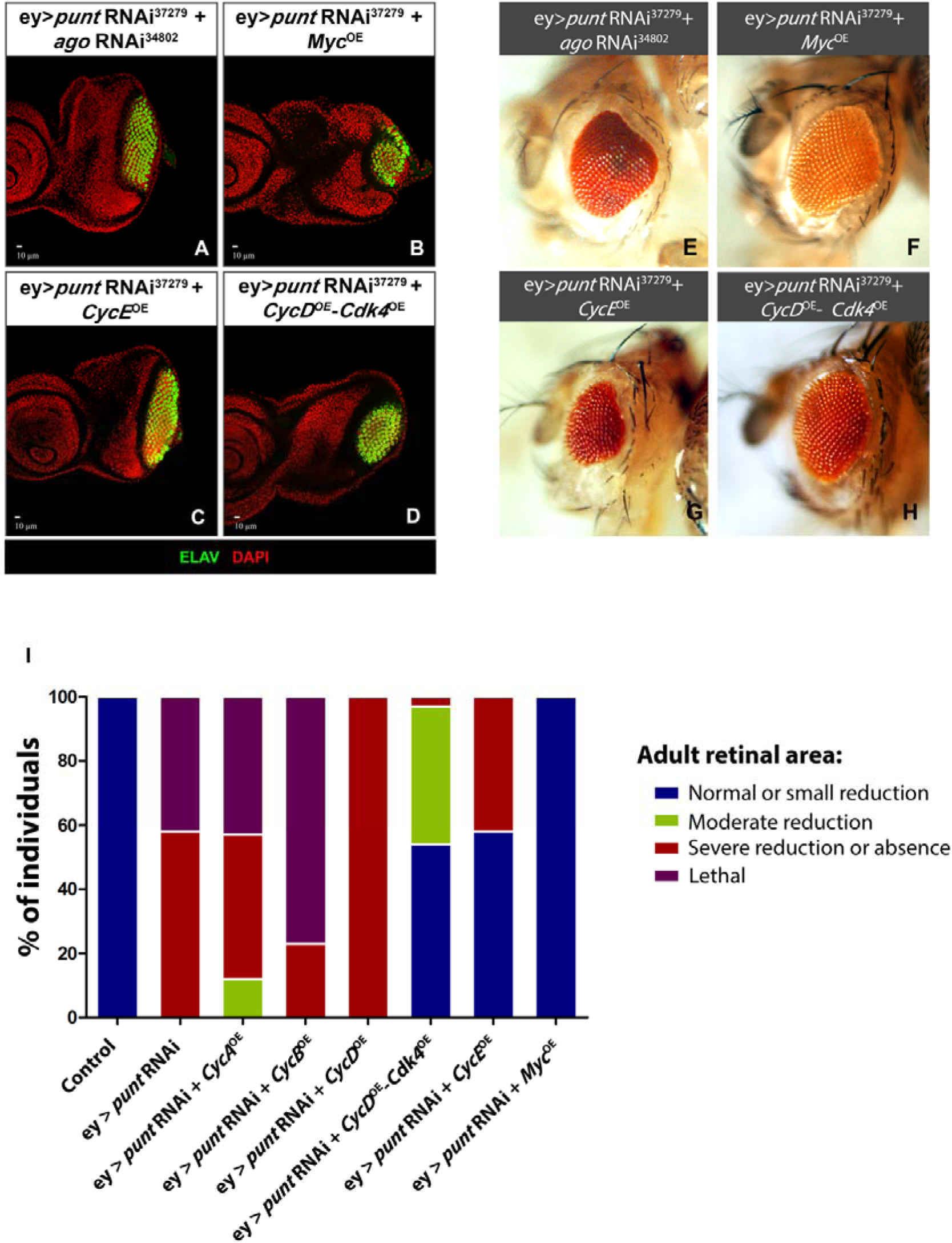
Overexpression of Myc, CycE, and CycD-Cdk4 rescue initiation of photoreceptor differentiation caused by Punt depletion. (A-I) In a similar manner to *ago* RNAi (A, E), overexpression of *Myc* (B, F, I), *CycE* (C, G, I) and *CycD-Cdk4* (D, H, I) recover the initiation of photoreceptor differentiation in eye discs (A-D) and retinal formation (E-H) in the ey>*punt* RNAi^37279^ genetic background. (A-D) Eye imaginal discs of the indicated genotypes stained with DAPI (DNA, red) and anti-ELAV (photoreceptors, green). Scale bars correspond to 10 μm. (M) Percentage of individuals of the indicated genotypes presenting normal adult retinal area or small reduction of adult retinal area (+++), moderate reduction of adult retinal area (++), severe reduction or absence of adult retinal area (+ or no retina) and lethality in pupa (n=43−96).

### 3.4. CtBP is a negative regulator of Mad activation by phosphorylation

*Drosophila* CtBP was initially reported as a transcriptional co-repressor able to form complexes with other DNA-binding transcription factors, such as Hairy and Eyeless, to suppress transcription of their target genes (Bianchi-Frias et al., 2004; Hoang et al., 2010; Nibu et al., 1998; Poortinga et al., 1998). Interestingly, *hairy* was shown to be a Dpp target expressed ahead of the MF, where it is proposed to contribute to the pace of furrow movement by restricting expression of atonal, a pro-neural transcription factor (Brown et al., 1995; Greenwood and Struhl, 1999; Spratford and Kumar, 2013). However, a different study could not identify any specific role for Hairy in the regulation of the MF (Bhattacharya and Baker, 2012). Furthermore, CtBP mutant adult retinas were reported to contain more ommatidia than wild-type (Hoang et al., 2010), and CtBP was described to interact with the transcription factor Danr, which contains a PXDLS motif and plays a role in specification and patterning through the regulation of atonal (Curtiss et al., 2007). As we showed above, CtBP works as a negative regulator of Dpp signaling in the eye disc. Thus, we aimed to distinguish whether CtBP works downstream of Mad activation working together with transcription factors regulated by phosphorylated Mad (pMad), like Hairy, or at the level of the Dpp pathway itself, upstream of Mad phosphorylation. For that aim we generated eye discs mostly composed of loss-of-function CtBP mutant cells, by early and extensive induction of mitotic CtBP mutant clones (*CtBP*^KG07519^) using eyeless-flippase (eyflip>CtBP^KG07519^) (Figure S6 and Figure 4). Strong downregulation of CtBP expression is observed in these eye discs (Figure S6), and we showed that *CtBP*^KG07519^ genetically interacts with *punt* RNAi loss-of-function (Figure S7) validating the interaction identified with the UAS-RNAi lines. Importantly, in *CtBP* mutant discs, we observed a strong upregulation of pMad (Figure 4D, 4D’, 4E and 4E’), with increased intensity and extensive broadening anterior to the differentiating cells when compared with control eye discs. As expected, overexpressing *punt* also caused pMad upregulation (Figure 4D, 4D’, 4F and 4F’), albeit weaker, which can be attributed to a wider progression of the MF and a sustained downregulation of Mad activation in differentiated cells posterior to the MF. In both genotypes, ey>*punt*^OE^ and eyflip>*CtBP*^KG07519^, retinal patterning was significantly affected (Figure 4A-4C). Remarkably, the overexpression of *CtBP* anterior to the MF, under control of the optix-Gal4 driver, was sufficient to strongly downregulate Mad activation (Figure 5), without inhibiting Mad protein expression (Figure S8). As CtBP appears to be sufficient for inhibition of Mad activation, we tested if upregulation of CtBP expression levels could contribute to the absence of retinal differentiation in *punt* loss-of-function. However, we could not detect significant changes in CtBP expression when *punt* RNAi was induced using ey-Gal4 or in mitotic clones (Figure 6). Overall, these results show that CtBP is a negative regulator of Dpp signaling in the eye disc acting upstream of Mad activation by phosphorylation.

**Figure 4.**
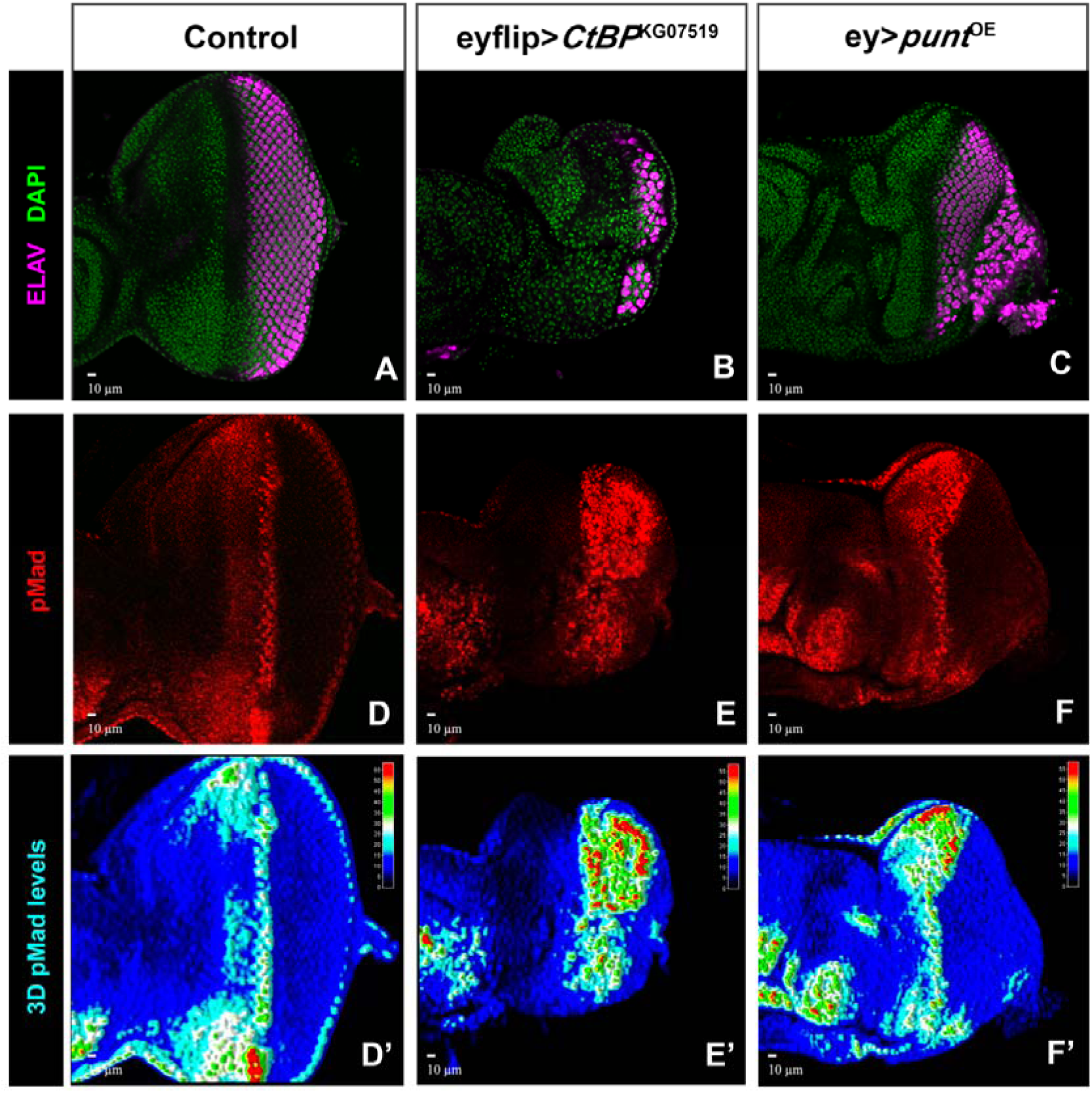
CtBP knockdown upregulates Mad activation by phosphorylation. (A-E’) *CtBP*^KG07519^ mutant eye discs were generated by eyeless-flippase induction (eyflip>CtBP^KG07519^). A broad and intense pattern of Mad activation (pMad) is detected (B, E, E’). The induction of Dpp signaling by ey>*punt*^OE^ leads to a precocious differentiation of the eye imaginal disc and pMad detection in regions anterior to the MF (C, F, F’). (D’, E’, F’) 3D histograms of pMad patterns in eye discs of the indicated genotypes. Eye imaginal discs of the indicated genotypes stained with DAPI (DNA, green), anti-ELAV (photoreceptors, magenta) and anti-pMad (red). Scale bars correspond to 10 μm.

**Figure 5.**
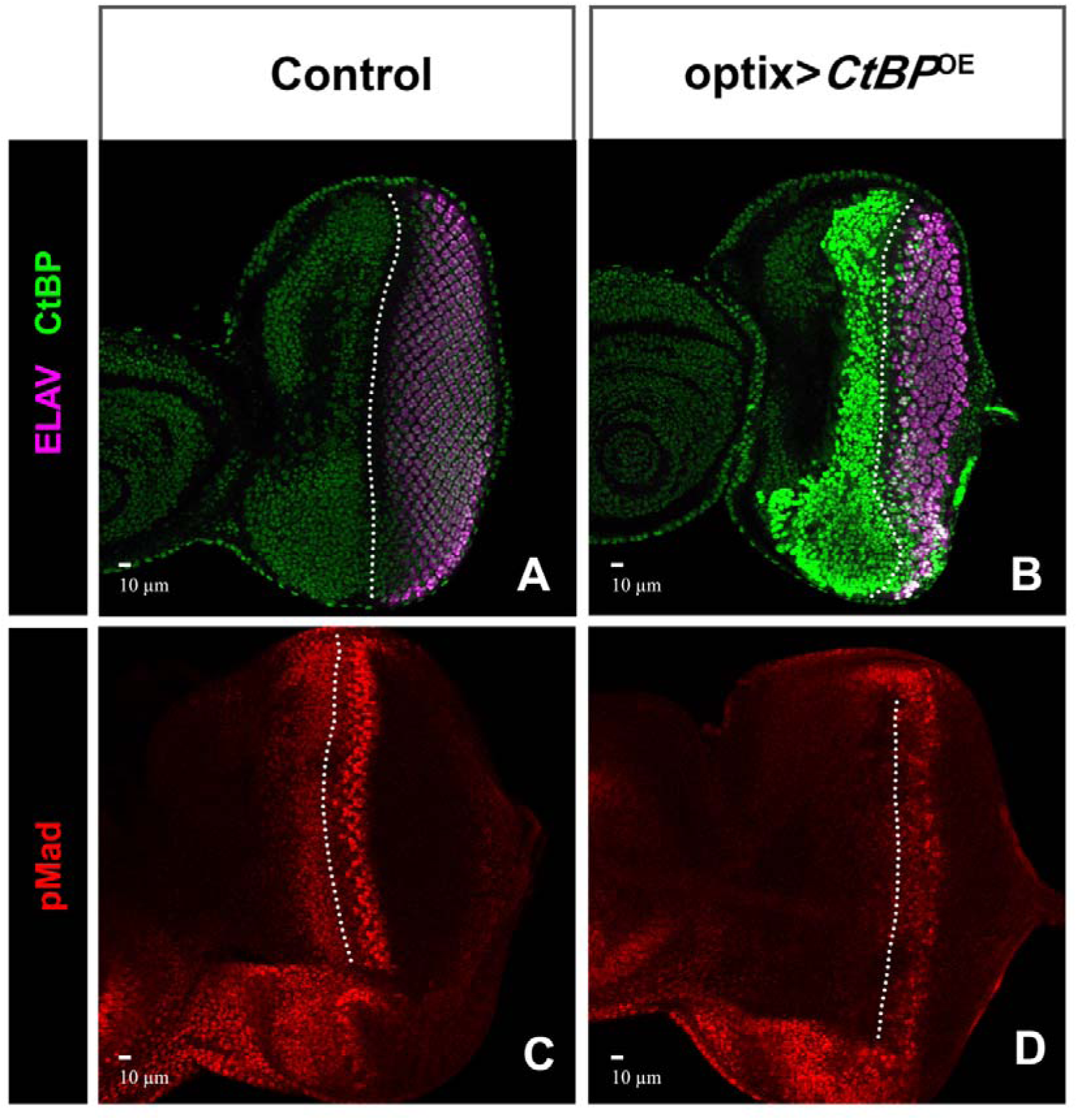
CtBP inhibits Mad activation. (A-D) Eye imaginal discs of *optix*-Gal4 (control) (A and C) and *optix*>*CtBP*^OE^ (B and D) stained with anti-CtBP (green), anti-ELAV (photoreceptors, magenta), and anti-pMad (red). Scale bars correspond to 10 μm. The dashed line marks the morphogenetic furrow (MF).

**Figure 6.**
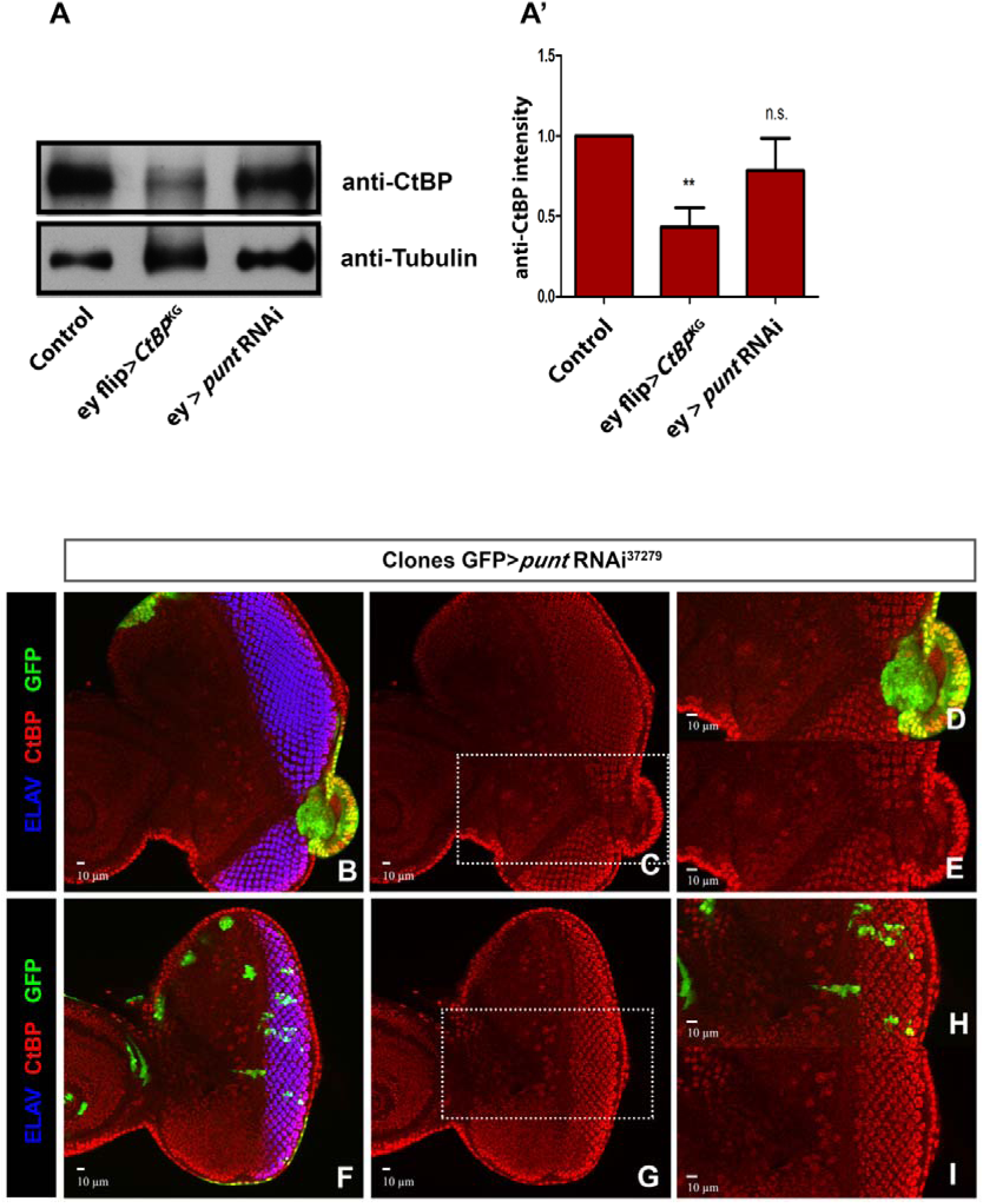
The knockdown of Punt does not alter CtBP levels. (A) Immunoblotting analysis of CtBP in control, eyflip> *CtBFP*^KG075l9^ and ey>*punt* RNAi^37279^ imaginal eye discs lysates. CtBP expression is decreased in *CtBP*^KG07519^ mutant eye discs, however ey>*punt* RNAi^37279^ have similar CtBP protein levels to the control. (A’) CtBP band intensities (relative to control) were quantified and the mean values are presented in a bar graph (n = 3). Data are normalized to the levels of control (n.s. means no statistical difference between samples; **p < 0.01; error bars represent SEM). (B-I) *punt* RNAi^37279^ clones were induced in the *Drosophila* eye disc at 48 hours (B, C, D, E) and 72 hours (F, G, H, I) after egg laying and analyzed at the wandering L3 stage. No alterations in CtBP expression are observed. Clones are marked positively by the presence of GFP (green). The imaginal eye discs were stained with anti-CtBP (red) and anti-ELAV (photoreceptors, blue). D-E and H-I show magnifications of the inset shown in C and G, respectively.

### 3.5. CtBP cooperates with Dad for inhibition of Mad activation

An analysis of the interaction of *ago* and *CtBP* with the I-Smad *dad* (Kamiya et al., 2008; Tsuneizumi et al., 1997) revealed additional details on the role of both genes in Dpp signaling during eye disc growth and patterning. Overexpression of Dad inhibits differentiation in imaginal eye discs, as well as in adult eyes (Figure 7B and E), resembling the phenotype caused by *punt* loss-of-function. However, the simultaneous overexpression of *dad* with *CtBP* RNAi (Figure 7C and F) led to a partial recuperation of photoreceptor differentiation. Control imaginal eye discs (Figure 7D’) showed a sharp and intense pMad band close to the MF and a broader less intense anterior domain (Firth et al., 2010; Vrailas and Moses, 2006). As expected, in eye imaginal discs overexpressing *dad*, pMad was reduced to residual levels (Figure 7E’, Figure S9), supporting the knockdown of Dpp-BMP2/4 signaling pathway by this I-Smad (Kamiya et al., 2008; Tsuneizumi et al., 1997). However, simultaneous overexpression of *dad* with *CtBP* RNAi partially rescued Mad activation in and anterior to MF (Figure 7F’). Furthermore, we observed that simultaneous overexpression of both Dad and CtBP in the anterior domain enhanced the inhibition of Mad activation (Figure S10). These results suggest that Dad requires the function of CtBP for efficient inhibition of Mad phosphorylation and that CtBP acts as a negative regulator of Dpp-BMP2/4 signaling pathway, possibly in parallel and/or downstream of Dad.

**Figure 7.**
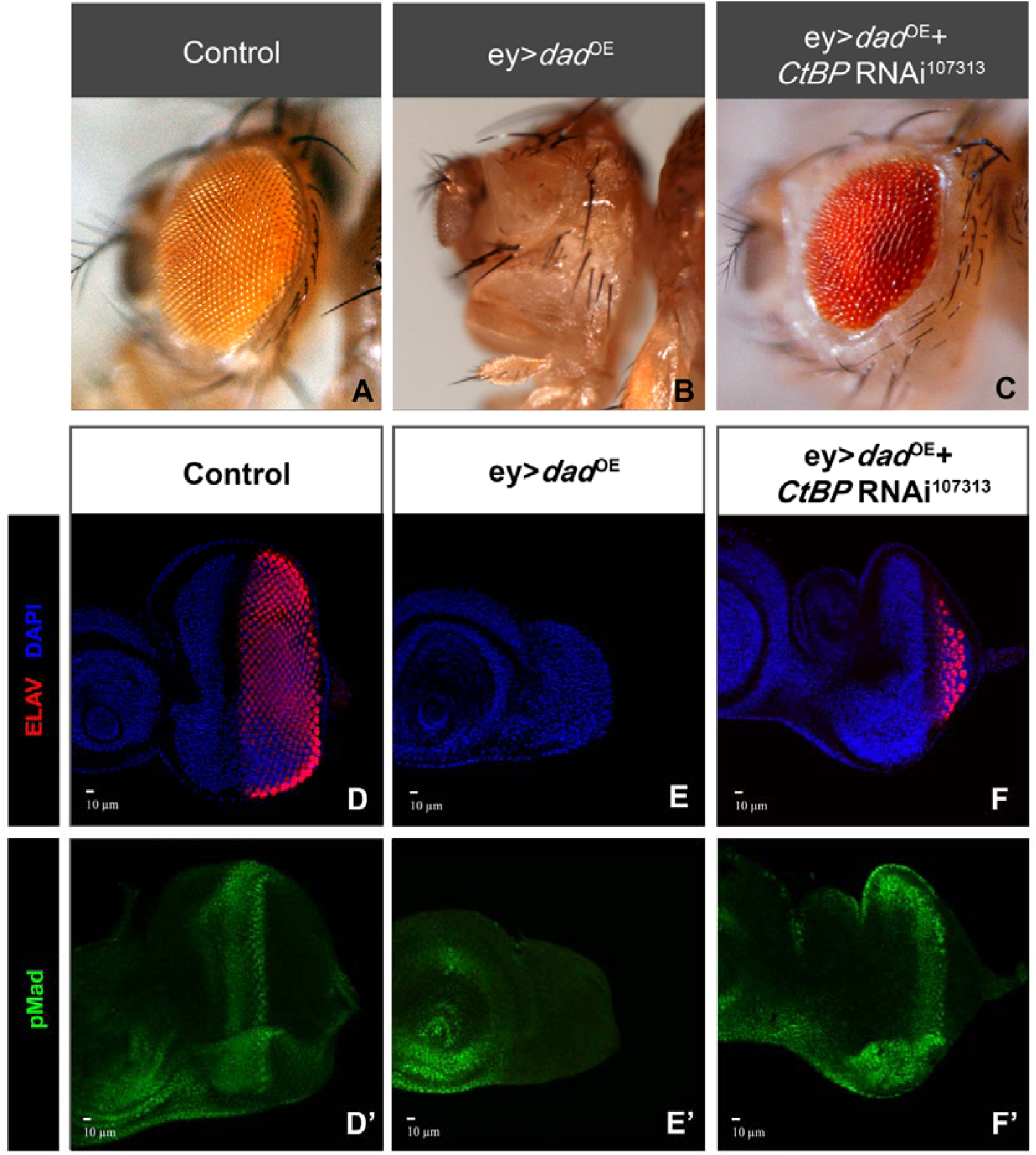
CtBP is required for Dad-mediated downregulation of Mad activation. (A-C) ey>*dad*^OE^ shows a very strong eye phenotype without retinal formation (B).However, retinal differentiation is rescued by co-expression of *CtBP* RNAi^107313^ (D-F’) The downregulation of Dpp signaling using ey>*dad*^OE^ inhibits photoreceptor differentiation in the eye disc (E). However, the co-expression of RNAi for *CtBP* together with overexpression of *dad* (ey> *dad*^OE^+*CtBP* RNAi^107313^) partially rescued differentiation (F). In ey>*dad*^OE^ eye discs, pMad is reduced to low basal levels (E’).Overexpression of Dad together with RNAi for *CtBP* rescued Mad activation (F’). Eye imaginal discs of the indicated genotypes were stained with DAPI (DNA, blue), anti-ELAV (photoreceptors, red), and anti-pMad (green). Scale bars correspond to 10 μm.

## 4. Discussion

The development of the *Drosophila* eye has served as a model system to study tissue patterning and cell-cell communication. Several key signaling pathways are conserved from flies to vertebrates, including TGFβ, Hh, Wg/Wnt, Notch, EGFR and JAK/STAT pathways (Zecca, Basler et al. 1995, Lee and Treisman 2001, Bach, Vincent et al. 2003, Reynolds-Kenneally and Mlodzik 2005). Dpp-BMP2/4 signaling plays an essential role in *Drosophila* development but we still have an incomplete knowledge of the regulation and functions of the Dpp-BMP2/4 during eye differentiation. Therefore, in this work we have taken advantage of an eye-targeted combinatorial screen to analyze the contribution of new Dpp-BMP2/4 genetic interactors. We studied a set of 251 genes with identified or putative functions in eye development (Marinho et al., 2013), and we identified four genes whose knockdown was able to significantly rescue eye-targeted loss-of-function for *punt* receptor: *brk*, *dad*, *ago* and *CtBP*. Co-induction of RNAi for each of these hits was able to rescue absence of retinal differentiation caused by *punt* RNAi expression (Figure 1) suggesting that these four candidate genes act as negative modulators of Dpp-BMP2/4 signaling in the eye disc.

The transcription factor Brk is a negative repressor of Dpp signaling which competes with activated Mad, blocking the stimulation of Dpp target genes (Bray, 1999; Campbell and Tomlinson, 1999; Jazwmska et al., 1999; Saller and Bienz, 2001). In the eye imaginal disc, Brk expression is detected at the very anterior region of the disc (Firth et al., 2010), and clonal *brk*-overexpression blocks the onset of the MF when clones are positioned along the disc margins (Baonza and Freeman, 2001), which represents a Dpp signaling loss-of-function phenotype. In the wing disc, the mechanisms by which Dpp controls patterning and growth have been intensively studied (Akiyama and Gibson, 2015; Barrio and Milan, 2017; Martin et al., 2017; Matsuda and Affolter, 2017; Sanchez Bosch et al., 2017). Dpp is expressed along the Anterior/Posterior boundary and the resulting gradient is required to establish distinct expression domains for targets (including *salm* and *omb*) involved in patterning.

However, formation of a Dpp gradient is not required for its ability to repress *brk* transcription and promote tissue growth (Sanchez Bosch et al., 2017). Here, we demonstrate that also in the eye disc, the requirement for Punt function, and therefore Dpp signaling, in growth and retinal differentiation can be bypassed by removing Brk repression, leading to a differentiated eye.

We also identified Ago, the *Drosophila* orthologous of Fbw7 in mammals and the F-box specificity subunit of the SCF-Ago E3 ubiquitin ligase, as a negative regulator of Dpp signaling. Ago protein is involved in cell growth inhibition by ubiquitination of several proteins, such as Myc and CycE (Moberg et al., 2001; Moberg et al., 2004). Loss of Ago in imaginal discs causes an accumulation of CycE and Myc, which drive cell growth and proliferation (Moberg et al., 2004). The rescue of retinal differentiation induced by *ago* knockdown in a *punt* RNAi background was mimicked by overexpression of Myc. Interestingly, Myc was identified as a target of Brk repression in the wing disc, where it was proposed that Dpp signaling inhibits Brk, thereby inducing expression of Myc that contributes significantly to Dpp-stimulated tissue growth (Doumpas et al., 2013). Thus, both Brk and Ago functions converge on the downregulation of Myc expression, at the transcriptional and post-transcriptional level, respectively. This supports the hypothesis that the mechanism for their genetic interaction with Punt that we identified in this study includes the contribution of Myc upregulation and tissue growth. We have also recently demonstrated that overexpression of Myc and Punt is able to enhance tissue growth and retinal differentiation in the eye disc (Martins et al., 2017). In here, we also show that other growth-stimulating conditions like overexpression of CycE or CycD-Cdk4 (Datar et al., 2000a) can partially rescue Punt knockdown. Overall, our results suggest that in the eye discs with reduced Dpp signaling, promoting tissue growth is sufficient to create the conditions required for the Dpp-dependent initiation of photoreceptor differentiation at the disc margins.

A third Dpp negative regulator identified in here was the I-Smad, Dad. In mammals the Dad orthologous, Smad 6 and 7, downregulate TGFβ signaling pathway by competing with R-Smads for receptors or for co-Smad interactions and also by targeting the receptors for degradation (Miyazawa and Miyazono, 2016). In *Drosophila*, Dad stably associates with Tkv receptor and thereby inhibits Tkv-induced Mad phosphorylation and nuclear translocation (Inoue et al., 1998; Kamiya et al., 2008). In the eye disc, overexpression of Dad in the Dpp-expression domain was shown to block the MF at the disc margins (Niwa et al., 2004), and pMad levels are upregulated in *dad*^212^ mutant clones (Ogiso et al., 2011). In here, we identify a significant role for Dad negative regulation of Dpp signaling in eye development, acting downstream of Punt receptor activity.

In this work we showed that knockdown of CtBP activity can compensate for a reduced level of Punt function, rescuing Dpp signaling to levels sufficient to restore retinal differentiation in a *punt* RNAi background. CtBP is a transcriptional repressor that functions as part of a complex containing enzymes that influence transcription by covalently modifying histones and influencing nucleosome packing and the binding of chromatin-associated proteins (Chen et al., 1999; Chinnadurai, 2002; Kim et al., 2005). Acting as a transcriptional co-repressor CtBP has been proposed to have both positive and negative contributions for Dpp/BMP signaling efficiency. On one hand, for a positive contribution, CtBP contributes to Shn/Mad/Med repression activity (Yao et al., 2008), which mediates Dpp-dependent Brk repression. However *brk* is not ectopically expressed in CtBP clones in the wing disc (Hasson et al., 2001) suggesting that CtBP is not essential for Dpp signaling activation in that tissue. On the other hand, in mammalian cells CtBP interacts with Smad6 to repress BMP-dependent transcription (negative input), a nuclear Smad6 role that is independent of its binding to receptors (Lin et al., 2003). Our results suggest that the interaction between CtBP and I-Smad could be conserved in *Drosophila*, as the Dpp repression by Dad requires the function of CtBP. Additionally, we showed that CtBP works upstream of Mad activation, in parallel or downstream to Dad. Interestingly, we could not identify CtBP-interaction motifs (PxDLS) in Dad, the *Drosophila* I-Smad, suggesting that distinct molecular mechanisms support the *CtBP-dad* genetic interaction and the negative regulation of Mad activation exerted by CtBP expression in the eye disc (Figure 5 and 6).

## Acknowledgements

We thank David Arnosti, the Bloomington Drosophila Stock Center, the Vienna Drosophila RNAi Center, the Undergraduate Research Consortium in Functional Genomics, the Drosophila Genetic Resource Center, and the Developmental Studies Hybridoma Bank for reagents; Paula Sampaio (ALMF, IBMC) for technical support. We also thank Eva Carvalho and Rita Pinto for excellent technical assistance during this study.

This work is a result of the project Norte-01-0145-FEDER-000008 - Porto Neurosciences and Neurologic Disease Research Initiative at I3S and the project Norte-01-0145-FEDER-000029 - Advancing Cancer Research: From basic knowledge to application, both supported by Norte Portugal Regional Operational Programme (NORTE 2020), under the PORTUGAL 2020 Partnership Agreement, through the European Regional Development Fund (ERDF). NE is supported by doctoral grant from FCT (SFRH/BD/95087/2013). PSP is a recipient of a Portuguese “Investigator FCT” contract. The funders had no role in study design, data collection and analysis, decision to publish, or preparation of the manuscript.

